# Culture Conditions Differentially Regulate the Inflammatory Niche and Cellular Phenotype of Tracheo-Bronchial Basal Stem Cells

**DOI:** 10.1101/2024.09.04.611264

**Authors:** Shubha Murthy, Denise A. Seabold, Lalit K. Gautam, Adrian M. Caceres, Rosemary Sease, Ben A. Calvert, Shana Busch, Aaron Neely, Crystal N. Marconett, Amy L. Ryan

**Author notes:** These authors contributed equally to this manuscript (Shubha Murthy, Crystal N. Marconett). Correspondence: *Amy L Ryan (*).

## Abstract

Human bronchial epithelial cells (HBECs) derived from the tracheo-bronchial regions of human airways provide an excellent *in vitro* model for studying pathological mechanisms and evaluating therapeutics in human airway cells. This cell population comprises a mixed population of basal cells (BCs), the predominant stem cell in airways capable of both self-renewal and functional differentiation. Despite their potential for regenerative medicine, BCs exhibit significant phenotypic variability in culture. To investigate how culture conditions influence BC phenotype and function, we expanded three independent BC isolates in three media, airway epithelial cell growth medium (AECGM), dual-SMAD inhibitor (DSI)-enriched AECGM, and Pneumacult Ex plus (PEx+). Extensive RNA sequencing, immune assays and electrical measurements revealed that PEx+ media significantly drove cell proliferation and a broad pro-inflammatory phenotype in BCs. In contrast, BCs expanded in AECGM, displayed increased expression of structural and extracellular matrix components at high passage. Whereas culture in AECGM increased expression of some cytokines at high passage, DSI suppressed inflammation altogether thus implicating TGF-β in BC inflammatory processes. Differentiation capacity declined with time in culture irrespective of expansion media except for PLUNC expressing secretory cells that were elevated at high passage in AECGM and PEx+ suggestive of an immune modulatory role of PLUNC in BCs. These findings underscore the profound impact of media conditions on inflammatory niche and function of *in vitro* expanded BCs. The broad pro-inflammatory phenotype driven by PEx+ media, in particular, should be considered in the development of cell-based models for airway diseases and therapeutic application.

**NEW & NOTEWORTHY:** Airway basal cells, vital for airway regeneration and potential therapies, show significant changes based on culture conditions. Our study reveals that media composition and culture duration greatly affect basal cell properties with profound changes in the pro-inflammatory phenotype and extracellular matrix deposition driven by changes in growth conditions. These results underscore the critical impact of culture conditions on BC phenotype, influencing cell-based models for airway disease research and therapy.

## INTRODUCTION

Human bronchial epithelial cells (HBECs), derived from the tracheo-bronchial regions, are the gold standard for studying airway pathology and evaluating therapeutics. This mixed population includes basal cells (BCs), the primary stem cells in the airways, known for their self-renewal and differentiation capabilities^1–5^. Due to their regenerative properties, BCs are considered a core cell type for airway regeneration and with potential for use in cellular therapy for respiratory disease. Unfortunately, the regenerative capacity of BCs exhausts because of aging and airway disease. Similar observations are made during *in vitro* culture of isolated cells^6, 7^. This significantly limits their use in high-throughput studies, expansion for cellular therapeutic approaches and investigating mechanisms regulating stem cell phenotype in airway disease and aging.

This study focuses on using *in vitro* expansion of airway basal cells, as a model of stem cell exhaustion, to understand the cellular mechanisms that impede long-term BC self-renewal both *in vitro* and *in vivo*. BCs are typically characterized by their expression of cytokeratin 5 (KRT5), tumor protein 63 (TP63), nerve factor growth receptor (NGFR), integrin alpha 6 (ITGA6 or CD49f) and podoplanin (PDPN)^3, 8, 9^. A subset of BCs (<20% in the murine airways) also express KRT14 and are also capable of responding acute lung injury^10^. More recently, in depth single cell RNA sequencing (scRNAseq) has identified sub-populations of BCs that have been segregated based on gene expression profiles^11–14^. This includes DLK1^+^ multipotent^15^, MUC4^+^/SCGB1A1^+^ secretory^16^ and FOXJ1^+^/DNAH5^+^ ciliated cells primed in the tracheobronchial regions of the airways^14^. However, the functional phenotype of these sub-populations is currently unknown.

Despite reflecting an *in vivo*-like functional phenotype *in vitro*, BCs undergo limited population doublings before losing their self-renewal and differentiation capabilities. Over the past decade, modifications to culture media have extended the number of passages over which BCs retain stem cell properties. For instance, SMAD signaling inhibition preserves BC proliferative capacity for up to 20 passages^7^. Commercial media like Pneumacult™ Ex+ further enhances BC proliferation compared to traditional media. Yet, BCs still rapidly undergo phenotypic changes, leading to stem cell exhaustion.

The effects of culture conditions on BC functions, such as extracellular matrix deposition and secretion of growth and pro-inflammatory factors, are not well understood^7, 17, 18^. Current *ex vivo* conditions may favor certain BC subpopulations, leading to a loss of stemness^7, 9, 10, 19^. To address this, our study evaluated the impact of three commercially available serum-free media, AECGM, DSI, and PEx+, on BCs using bulk RNAseq coupled with functional assays to analyze transcriptomic and phenotypic changes during *in vitro* aging. Our findings reveal that extended culture significantly alters BC phenotype, transcriptomic profile, and function, regardless of media conditions. Differentiation to ciliated, club, and goblet cells decreased with high passage, though media-specific effects were noted. AECGM and PEx+ media increased PLUNC expression, with PEx+ also promoting an inflammatory signature, while DSI suppressed inflammation despite some inflammatory gene enrichment. These results have important implications for using *ex vivo* expanded BCs in cell-based therapies and *in vitro* models for lung disease research.

## MATERIALS AND METHODS

### Isolation and culture of primary HBEC

Details of the donor lung tissues can be found in **Supplemental Table S1.** HBEC were isolated following previously published protocols^20^. In brief, HBEC isolation from human lung tissue from subjects with no prior history of chronic lung disease were performed with approval of IRB at USC (protocol #HS-18-00273). Briefly, proximal airways from the trachea, main stem bronchi and 2-3 further branching generations pf the cartilaginous airways were dissected into sections between 1-4 cm^2^ in size and placed in a solution (%w/v) of 0.1% Protease XIV (Sigma Aldrich, #P5147) and 0.001% DNase (Sigma Aldrich, #DN25) in DMEM/F12 (ThermoFisher, #11330032) overnight at 4°C. Using a #10 scalpel (Exel International #29550), epithelial tissue was gently scraped off, collected in DMEM/F12, and centrifuged at 400xg for 5 minutes. After red blood cell lysis in ACK lysis buffer (ThermoFisher #A1049201), epithelial cells were single cell dissociated by incubation with Accutase (Innovative Cell Technologies, #AT104) at 37°C for 30 minutes. Cells were then seeded at a density of 30×10^3^ cells/cm^2^ on PureCol (Advanced BioMatrix, #5005) coated tissue culture dishes in airway epithelial growth media (AECGM, Promocell, #C-21160) and passaged at 80% confluence. At each passage cells were seeded at 5×10^3^ cells/cm^2^. Cells were cultured in AECGM (Promocell) for one passage prior to changing into one of three medias at P1 including AECGM, AECGM enriched with dual SMAD inhibitors (DSI), A-83-01 (Selleckchem, #S7692 at 0.1µM) and Dorsomorphin (DMH1) (Selleckchem, #7146 at 0.1 µM) or PEX (Stem Cell Technologies, #05040) (**Supplemental Table S2**). Cells were analyzed at low passage (P1 to P3, donor dependent) and high passage (P4 to P6, donor dependent).

### Differentiation of HBEC at the Air-Liquid Interface (ALI)

HBEC were dissociated in TrypLE for 5 minutes at 37°C, collected by centrifugation at 300 x g for 5 minutes, resuspended in growth media and counted. 50,000 cells were plated on PureCol coated 24 well Transwells (Corning, #3470) and cultured for 2-3 days until reaching confluence. Confluent layers of HBEC were “air-lifted”, the apical media was removed, and the apical surface washed with 100 µl PBS. The basolateral media was changed from the respective growth media to differentiation media, Pneumacult ALI (Stem Cell Technologies, #05001) and cells were cultured for 28 days.

### Cilia beat frequency (CBF)

CBF measurements on ALI differentiated cells were performed as previously described^21^. Briefly, ciliary movements were captured using a 40x water immersion objective on an inverted Zeiss microscope system equipped with a heated (37°C)chamber and highspeed camera with video capture. CBF analyses were performed using Sisson-Ammons Video Analysis Software (SAVA) on at least 5 random fields with beating cilia per donor^22^. Results are expressed as CBF for whole field area and % area covered by actively beating cilia.

### Immunofluorescence (IF) staining

Cells were fixed with 4% paraformaldehyde (PFA) for 15 minutes at room temperature and washed with PBS. Cells were permeabilized using 0.5% Triton-X100 for 10 minutes and then blocked in PBS containing 5% normal donkey serum and 2 % BSA for 1-2 hours. All antibodies used for IF staining are listed in **Supplemental Table S3.** Cells were incubated in primary antibodies for 2 hours at room temperature, washed 3 x 5 minutes each in PBS and subsequently incubated in secondary antibodies for 1 hour at room temperature. Cells were mounted using ProLong Diamond Anti-Fade Mountant (Invitrogen #P36961) Images were collected with a Zeiss LSM 980 confocal microscope. Maximum intensity projections were processed using ImageJ software and images represented as single or merged.

### Immunoblotting

After 5 days of culture with AECGM, DSI or PEx+, BC monolayers were washed with PBS and lysed with RIPA (Sigma #R0278) supplemented with protease (Roche #11836153001) and phosphatase inhibitors (Roche #04906845001). Cells were incubated with the lysis buffer for 30 minutes on ice and vigorously vortexed every 10 minutes. Lysates were clarified at 16,000 x g for 10 minutes. The supernatants containing clarified lysates were used for protein estimation and SDS/PAGE resolution. Total protein was determined by DC Lowry and MicroBCA assays. Equivalent amounts of total proteins were resolved on 4-12% gradient BIS-TRIS gels (Invitrogen #NP0321BOX) under reducing and denaturing conditions, transferred to PVDF membranes (Immobilon P #IPVH00010) and blocked for at least 1 hour with TRIS buffered saline containing 0.1% Tween-20 (TTBS) and 5% non-fat dry milk (NFDM) (blocking buffer). Membranes were incubated overnight at 4°C on a rocker with primary antibodies diluted as indicated in **Supplemental Table S3**. After washing 3×10minutes with TTBS membranes were incubated with appropriate secondary antibodies for 1 hour at the dilutions indicated in **Supplemental Table S3**. After washing with TTBS, blots were developed with ECL Prime (Amersham) and imaged on AI600 imager (Amersham). Images were densitometrically scanned using ImageJ software. All proteins except for fibronectin 1 were normalized to ß-Actin. Fibronectin 1 was measured in conditioned media harvested from the 5-day culture. Media were centrifuged at 300xg for 10min to sediment cell debris and supernatants were frozen in aliquots at -80°C. Media equivalent to the same proportion of wells in which cells were cultured 1 were resolved on gradient gels as described above and immunoblotted with anti-fibronectin1 antibodies as indicated in **Supplemental Table S3**.

### Bulk RNAseq

Attached cells in 6 well plates at ∼80% confluence were washed 2x with PBS and RNA was isolated using Tri-reagent (ThermoFisher, #AM9738) on ice. 250 µl of Tri-reagent was added to 2 wells per condition and incubated on ice for 10 mins. Cell layer was scraped, and 2 technical replicate wells pooled to make 500 µl of RNA isolate per donor for 2 donors. RNA was prepped as described above and RNA was quantified using a µDrop plate, (ThermoFisher, #N12391) in the Varioskan plate reader. Absorbance was measured at A260, and purity assessed with the A260/A280 ratio. 10 µg of RNA underwent multiplexed poly-A purified library construction and subsequent next generation sequencing using an Illumina NovaSeq S2 at Azenta (formerly Genewiz, South Plainfield, NJ).

### Bioinformatics Analysis – Bulk Sequencing

Paired end FASTQ read files were assessed for quality using FastQC ^23^. Reads were cleaned by filtering out those with less than 90% of the read having a quality score of < Q30 and were subsequently aligned to the Homo.Sapiens.GRCh38.96 transcriptome using Kallisto (version 0.46.0) ^24^. To account for the wide range of raw sequencing depth between samples (35M to 135M), total read numbers were adjusted to ∼42M per sample [aligned read range: ∼31 - 45M] prior to calculation of read count per transcript (**Supplemental Table S4**). Kallisto-aligned pseudocount tables were filtered to include transcripts with counts per million > 50 (**Supplemental Table S4**) and were subsequently input into R (v 4.0.2 GUI 1.72 Catalina build 7847) and EdgeR was used to calculate differential expression and FDR correction of significance ^25^. Visualization of gene expression differences were performed by importing gene expression tables into R studio (v 1.4.1717) and utilizing the gplots package ^26^. Selection for top variable genes was done using the genefilter package ^27^ and visualized using the heatmap.2 function. Differentially expressed genes and FDR-corrected P values were input into Ingenuity Pathways Analysis (IPA) using all known direct and indirect relationships for genes with < 0.05 FDR-corrected p value. Radar plots for upstream regulatory enrichment were performed using the tidyverse ^28^, patchwork, and viridis packages in R studio^29^. Barplots and scatterplots were generated in R using base plotting functions.

### Electric Cell-substrate Impedance Sensing (ECIS)

The ECIS Z-Theta (Zθ) system (Applied Biophysics, Inc) was used to measure impedance, resistance, and capacitance of HBECs during their growth phase in culture. 60K HBECs were plated on 8-well disposable ECIS arrays (8W10E+, 40 electrodes/well, Applied Biophysics) and placed in the holder plate in a humid incubator at 37 °C and 5% CO2. Cell behavior was monitored over a 60-hour time frame over the entire frequency range (62.5 -64000 Hz). A minimum of 4 independent wells (n=4) per cell donor (N=3) per media condition (AECGM, DSI and PEx+) were evaluated. Each array was coated with PureCol (Advanced Biomatrix #5005), stabilized, and inoculated with cells using 6400 µl per well of a 1.5 × 10^5^ cells/ml cell suspension. For this study, the default optimal frequencies were used: resistance (R) 4000 Hz, impedance (Z) 32,000 Hz, and capacitance (C) 64,000 Hz.

### Statistical Analysis: Statistical Analysis

All data are presented as mean ± S.E.M. Statistical analysis was performed using one-way ANOVA followed by a Sidak’s multiple comparison test with a value of *p*<0.05 being considered statistically significant. Unpaired t-test was used to compare differences in ITGB6, TNF-α, and IL-13 expression. Unless otherwise stated the data is collected from a minimum of 3 experimental replicates (n=3) and a minimum of 2 biological donors (RNA-seq, N=2) or 3 biological donors (all other experiments, N=3). Bulk RNAseq was fitted to a negative binomial model in EdgeR and testing performed using the QL-F test for small numbers of replicates. FDR correction for raw p values was performed using P. Adjust in R. Pathways enrichment was performed using IPA using the Fisher’s Exact Test with a Benjamini-Hochberg FDR correction.

## RESULTS

### Culture-dependent changes in BC phenotype with time in culture

To evaluate the impact of culture media on BC phenotype primary HBEC, isolated from a minimum of 3 donor lungs, were cultured in AECGM for one passage prior to sequential culture in three independent medias as shown in the diagram in **Figure 1A**. An immediate and notable impact of media on HBEC phenotype is shown in the phase contrast images in **Figure 1B**. Both AECGM and DSI^7^ maintain a similar cuboidal phenotype, whereas culture in PEx+^30–32^ leads to a more elongated and clustered growth of HBEC. Changes in the expression key BC markers over time highlight changes in KRT5 expression with levels higher in low passage with DSI and PEx+, decreasing over time and vice versa in AECGM with KRT5 levels low and rising over time in culture **(Fig. 1C).** P63 is a key marker of airway basal stem cells, and its expression and function is dependent on alternative promoter usage and/or splicing events that generate several different isoforms. TA-p63 and ΔN-p63 are two major isoform groups representing the transactivating and repressive isoforms, respectively whose transcription is driven from separate promoters. The ΔN-isoform lacks the functional transactivating domain and is the predominant isoform associated with basal cell stemness^33^. We, therefore, evaluated ΔN-p63 expression in BCs at low and high passage in AECGM, DSI or PEx+ and found a reduction in positive cells particularly in PEx+ and AECGM with DSI generating brighter cells and retaining expression in most cells in the culture **(Fig. 1D).**

**Figure 1:**
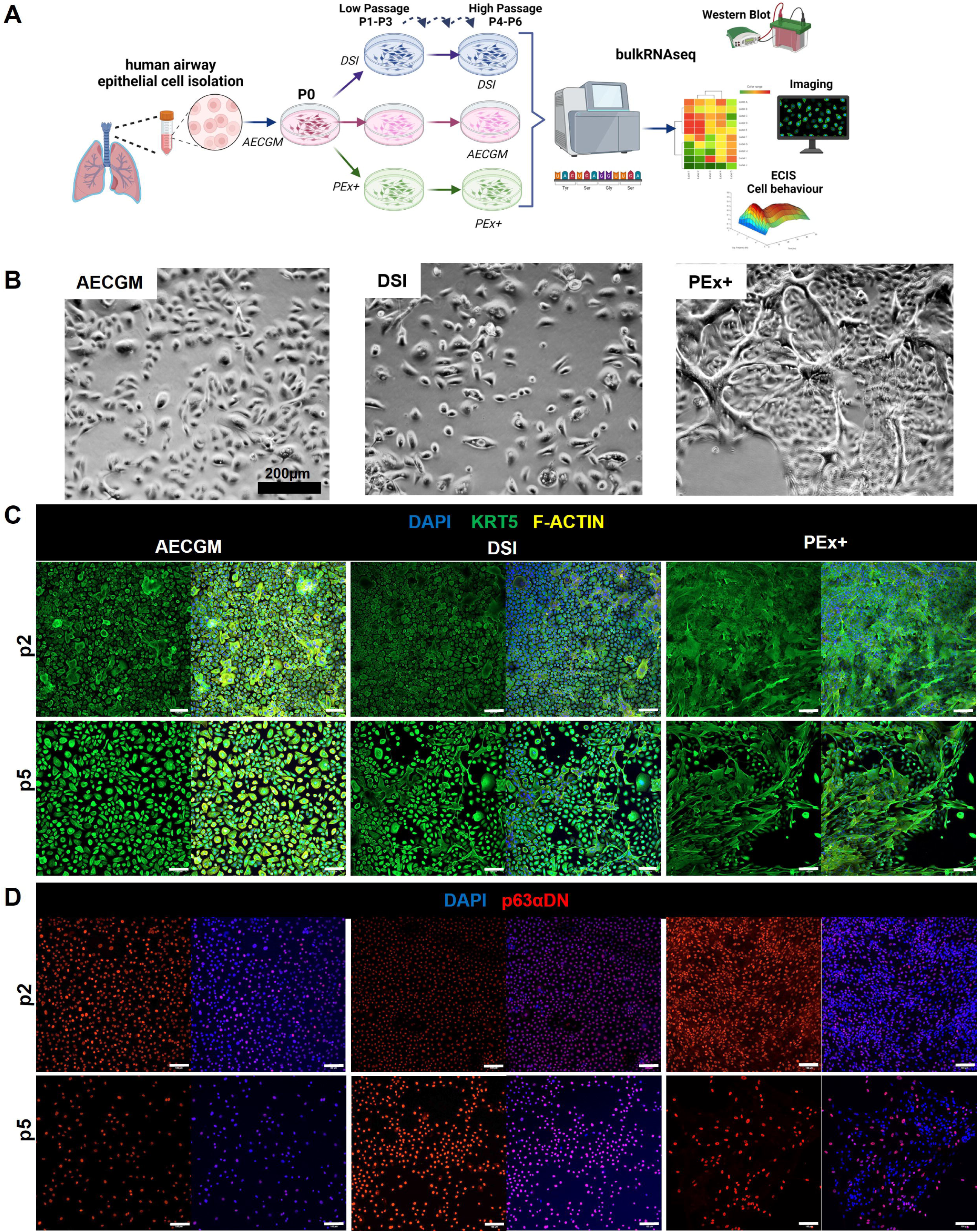
Long-term Culture Alters BC Phenotype. **(A)** Schematic of isolation, expansion, and subculturing of BCs in AECGM, DSI or PEx+ media. Cells were harvested at either low (p1-p3) or high passage (p5-p6) for bulk RNA seq, immunofluorescent (IF) detection of stem cell markers, Western blotting (WB), or growth kinetics using ECIS. In addition, low and high passage BCs cultured in the 3 different media, were differentiated for 28 days at the air-liquid-interphase (ALI) to assess differentiation to ciliated, secretory or goblet cells and ciliary beat frequency. This schematic was made with Biorender.com. **(B)** Phase contrast images of low passage BCs cultured for 5 days in the 3 different media. Scale bar=200µm. Representative IF images from single donor showing KRT5 (green) and KRT5-F-ACTIN (yellow)-DAPI (blue) merged expression **(C)** and P63αΔN (red) and P63αΔN-nuclei (DAPI, blue) merged images **(D)** at low and high passage in AECGM, DSI or PEx+. Scale bar=100µm.

ALI-induced differentiation of the expanded BCs was compared after 28 days of differentiation (**Fig. 2**). BCs were first expanded in the three media and then differentiated at low or high passage in in Pneumacult ALI media to induce differentiation equally over all cultures. We recently mapped the structure to function relationships in human airways and Pneumacult ALI best matched the differentiation model for larger branching generations in the human airways^34^. Differentiation capacity is less dependent upon expansion media than it is on differentiation media^34, 35^. We observe that the capacity for differentiation does substantially and rapidly decrease with time in culture irrespective of the expansion media (**Fig. 2**). IF images from single donor are shown comparing all three expansion medias at low (**Fig. 2A**) and high (**Fig. 2B**) passages exposed to AECGM, DSI or PEx+. The representative images highlight changes in the distribution of acetylated alpha tubulin (ATUB) expressing multiciliated cells, club cell 10Kda protein (CC10) expressing club cells, S-PLUNC expressing secretory cells, mucin5AC (MUC5AC) expressing goblet cells, and F-actin depicting cell junctions. Minimal media-dependent differences in low passage cells were observed. At low passage, cells expanded in all media maintained a tight cobblestone morphology with F-Actin-stained tight junctions defining cell boundaries. At higher passages there was a notable loss in differentiation with distinct differences in each media. AECGM cells did not generate multiciliated cells and to some extent retained intracellular MUC5AC and CC10 expression. A substantial change in cell morphology noted by an increase in size and flattening of the epithelium was observed with erratic tight junction formation and gaps in the culture as evidenced by F-actin immunostaining. Marked increase in S-PLUNC secretory cell marker expression was observed in AECGM. The addition of DSI to the media blocked differentiation at high passage to multiciliated, club, PLUNC^+^, secretory and MUC5AC^+^ goblet cell differentiation. Cells expanded in DSI had a larger and flatter cell morphology with a lack of tight junction formation. Expansion in PEx+ medium resulted in a similar lack of multiciliated cells, but this was associated with a notable decrease in expression of CC10 and MUC5AC club and goblet cell markers, respectively. Similar to AECGM, PEx+ expanded BCs demonstrated marked increase in S-PLUNC^+^ secretory cells at high passage and demonstrated a lack of organized tight junction arrangements in the epithelial sheet. Raw images from a minimum of 3 donors are included in the online data supplement for Figure 2, supporting the representative examples shown. While donor to donor variability is evident the same trends in differentiation capacity are observed in paired data. Donor #3 however, differed from the other two donors in that very few cells after expansion in AECGM remained on the filter after 28 days of differentiation making it difficult to appreciate much immunostaining. Functional properties of ciliated cells at low and high passages in the 3 different media were quantified and compared through analysis of ciliary beat frequency (CBF, **Fig. 2C**) and distribution of actively beating ciliated cells (% active area, **Fig. 2D**). Media-dependent effects on CBF at low passage was donor-to-donor-dependent. While CBF significantly increased in DSI and PEx+ for two donors, the third donor did not demonstrate any change resulting in no significant difference overall between media.. Extended passage significantly decreased active area of beating ciliated cells irrespective of the expansion media used except for donor #2 (**Fig2D**). Cilia coverage in donor #2 remained elevated when expanded in DSI but not AECGM or PEx+ media. This variability is likely due to the number of active stem cell clones in each passage of the cells rather than a specific impact of media on cell function.

**Figure 2:**
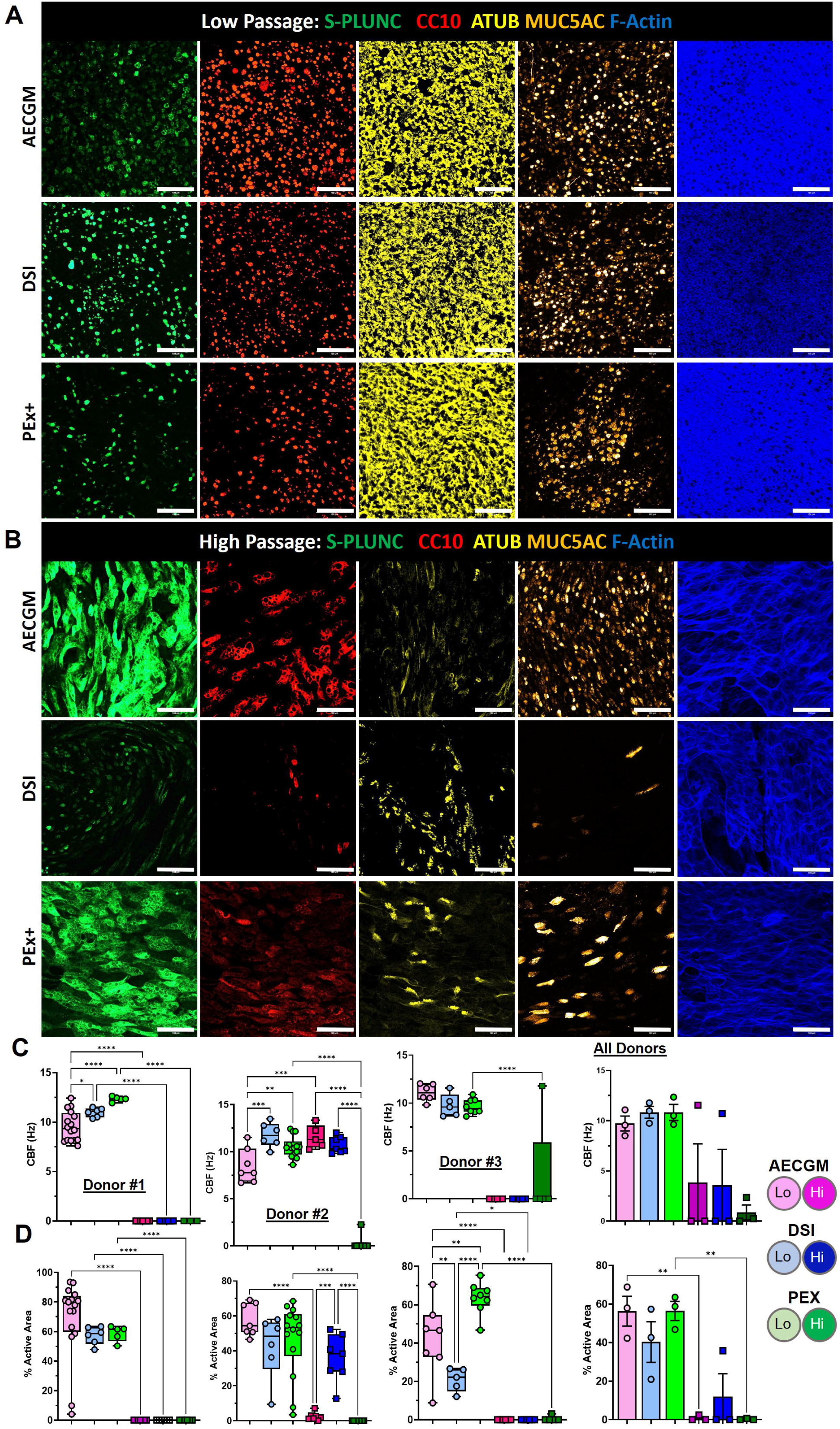
Culture Conditions Significantly Impact BC Differentiation. Representative ALI IF images from single donor at low **(A)** and high passage **(B)** expanded in AECGM, DSI or PEx+ and differentiated in Pneumacult ALI medium. Acetylated Tubulin (ATUB), CC10, PLUNC, mucin 5AC (MUC5AC) and F-actin expression are shown. Scale bar=100µm. **(C)** Functional properties of ciliated cells at low and high passages in the 3 different media were measured by ciliary beat frequency (CBF) and distribution of actively beating ciliated cells (% Active area). Mean±SEM of 3 donors, * p <0.05, **p<0.01, ***p<0.001, ****p<0.0001.

### Transcriptomic changes in primary HBEC occur during culture

Our research has identified significant transcriptomic changes in primary HBECs during culture. To investigate the impact of culture conditions on the transcriptomic profile of BCs in culture, we performed bulk RNA sequencing of BCs comparing low and high passage cells from two independent human lung donors (details in **Supplemental Table S1**). We observed that transcriptomic variability, which was noticeable at P0 ( **Supplemental Figure S1**), decreased after passaging, with gene signatures clustering most closely at the same passage (**Supplemental Figure S1** and **Fig. 3A-B**). Interestingly, closer clustering of gene signatures were observed between the following groups: DSI and PEx+ at low passage; and all medias at high passage (**Fig. 3A-B**). Four distinct clusters of differentially expressed genes (DEGs) emerged from our analysis: 1) Genes higher in AECGM at both low and high passages compared to the other two media; 2) Genes activated in P4 DSI and PEx+ compared to AECGM; 3) Genes showing donor-specific differences over time in response to DSI and PEx+ and 4) Genes that were downregulated with passage in all media (**Fig. 3B**). **Figure 3C** presents a three-dimensional plot illustrating the distinct gene expression changes between high (+) and low passage (-) across all three media types. A closer examination of cluster 4, which comprises genes downregulated with passage in all media, revealed that these genes were predominantly associated with a decrease in cell cycle-related gene expression (**Fig. 3C-F**). Specifically, genes like TOP2A, CENPF, CDC20, and MIK67 were downregulated at the transcriptomic level (**Fig. 3D**), suggesting a stalling of the metaphase to anaphase transition and subsequent mitotic exit from the cell cycle (**Fig. 3F**). The dot plot in **Figure 3G** further highlights that cell cycle-related genes are among the most significantly downregulated genes with time in culture. In addition to the decline in cell cycle-related gene expression, we observed a significant increase in DNA damage and repair pathways and Sirtuin signaling pathways over time, regardless of the culture medium (**Fig. 3E**). The decrease in cell proliferation was validated by quantifying protein expression of CENPA via western blot (**Fig. 3H**). CENPA expression decreased at high passage when expanded in AECGM or PEx+. However, expansion in DSI demonstrated some donor variability with two out of three donors demonstrating decrease and one donor showing equivalent expression suggestive of a higher BC proliferative capacity in this donor.

**Figure 3.**
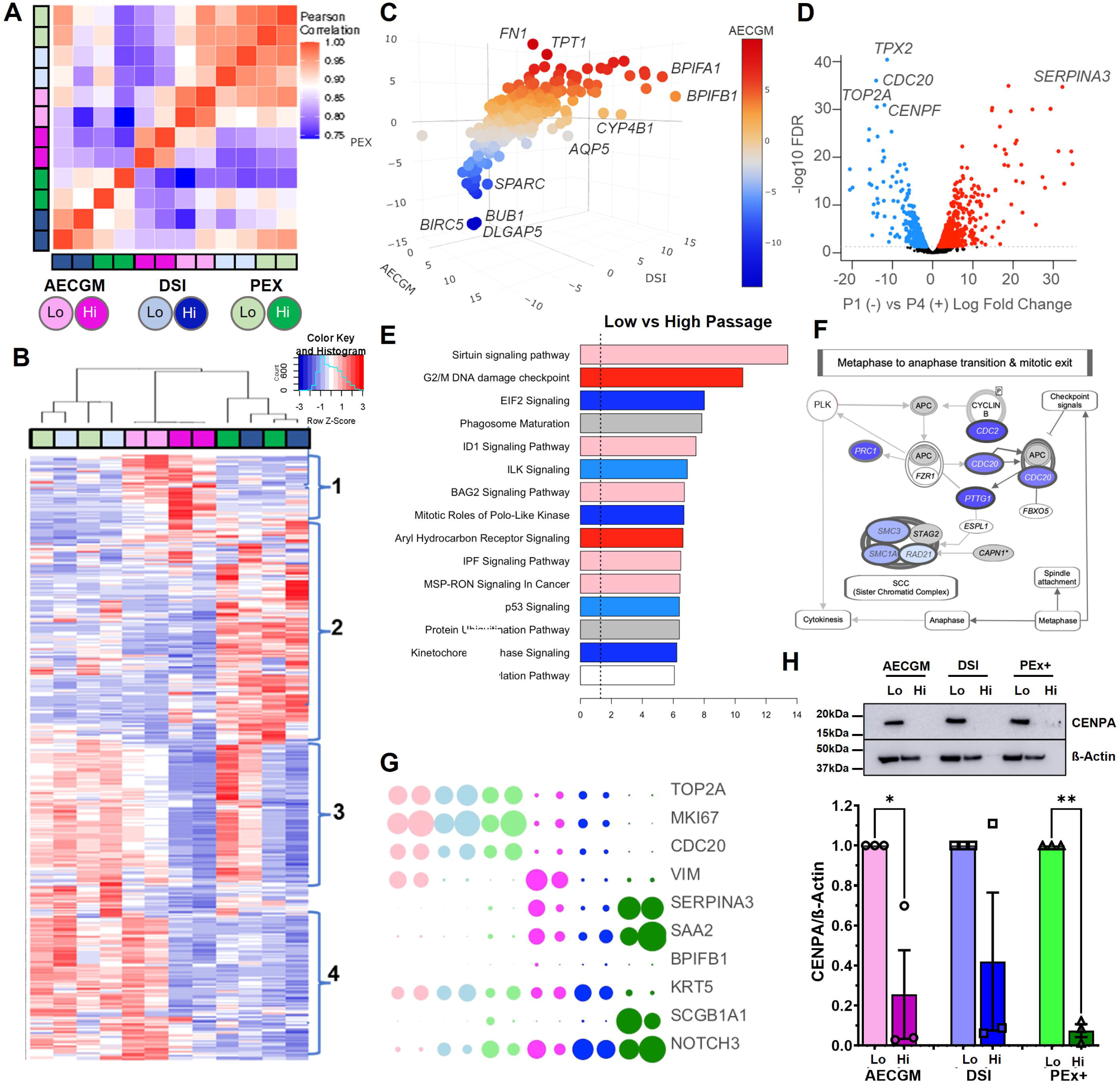
Culture In AECGM Media Generates Distinct Basal Cell Transcriptome Patterning. **(A)** Pearson correlation matrix of gene expression patterns across samples. (**B**) Unsupervised hierarchical clustering of top 20% of differentially expressed genes between high and low passage for all media types. (**C**) Three-dimensional plot showing distinct expression changes between all 3 media types. All axes represent log fold change in gene expression between high (+) and low (-) passage. Red = upregulated in AECGM at high passage, blue = downregulated in AECGM at high passage. (**D**) Volcano plot of differential expression between high (+) and low (-) passage for all media types. Red = increased expression, blue = decreased expression. (**E**) IPA pathways enrichment analysis for high (+) vs low (-) for all media types. Red (and shades of) = positive Z score of enrichment, blue (and shades of) = negative z score of enrichment, Grey = not able to calculate a z-score of activation. Data is plotted as - log10-BH corrected p value of pathway enrichment, the vertical dashed line indicates significance threshold. (**F**) Loss of metaphase separation observed in all media types. Canonical pathway shown. Blue = gene downregulated in high vs. low passage. (**G**) Dot plot of the top DEG between media and passage. (**H**) Representative immunoblots of CENPA and ß-Actin from single donor and quantitation of CENPA normalized to β-Actin in 3 donors. Mean±SEM. *p<0.05, **p<0.01.

IPA-enrichment analysis further confirmed significant negative enrichment of cell cycle-related pathways, including cell cycle replication, metaphase signaling, and polo kinase (Meiosis), along with positive enrichment of the G2/M checkpoint in all media (**Fig. 4**). While cell cycle-related pathways had the highest z-scores in both AECGM (**Fig. 4A**) and DSI (**Fig. 4B**), PEx+ (**Fig. 4C**) showed positive enrichment in Sirtuin signaling, cholesterol biosynthesis, and significant positive enrichment of NFR2 oxidative stress pathways, indicating an overrepresentation of metabolic regulatory pathways closely associated with culture in PEx+ (**Fig. 4**).

**Figure 4.**
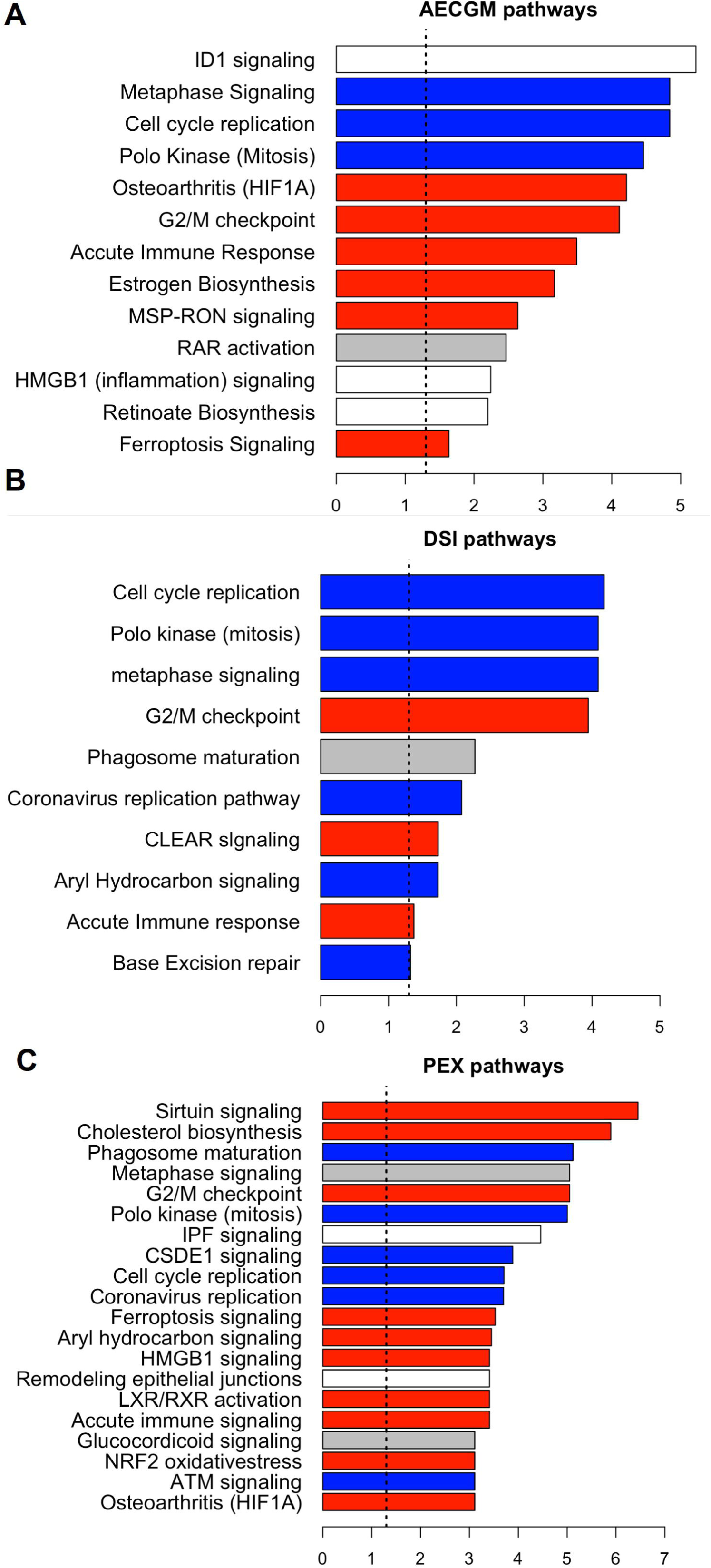
Media Specific Changes in IPA pathways associated with BC Expansion in Culture. IPA pathway enrichment analysis for high (+) and low (-) passage for AECGM **(A)**, DSI **(B)** and PEx+ **(C)**. Red = positive Z score of enrichment, blue = negative z score of enrichment, Grey = not able to calculate a z-score of activation and white has a z-score of activation of 0. Data is plotted as - log10-BH corrected p value of pathway enrichment, the vertical dashed line indicates significance threshold.

### ECIS measurements for real-time cellular behaviors

ECIS is a non-invasive technique used to monitor real-time cellular behaviors, including barrier function and cell migration. We, therefore, evaluated the impact of three different culture media on cellular behavior over time, using ECIS to measure key parameters such as barrier integrity and cell adhesion/spreading as reflected by resistance (R) and cell capacitance (C). Although a decrease in cell proliferation with passage was observed across all media (data not shown), distinct growth patterns emerged depending on the medium. To assess these differences, we seeded cells at a density of 75×10^3^ cells/cm² and tracked the electrical resistance of cell-covered electrodes across various frequencies (ranging from 62.5 Hz to 64,000 Hz) over a 60-hour period. The results, illustrated in three-dimensional plots, show significant media-dependent changes in resistance (**Fig. 5A**) and capacitance (**Fig. 5B**) within the first 10 hours post-seeding. These changes also depended on the frequency applied to the cultures. Resistance measurements taken at a low frequency provides insight into the barrier formation and integrity whereas capacitance measurements taken at high frequency are reflective of cell adhesion, spreading and confluence. For instance, at low frequencies (≤1 kHz), resistance increased rapidly in the first 5 hours for AECGM and the first 10 hours for PEx+, followed by a drop (**Fig. 5A, C-D**). After this decline, resistance then stabilized in PEx+, while in AECGM, it experienced a slower secondary increase before declining again. In contrast, resistance in DSI continued to rise over 48 hours until reaching its final value. At higher frequencies (>1 kHz), the initial increase in resistance during the first 10 hours was less pronounced. Taken together, these changes suggest media-dependent changes in rates at which the monolayers achieved barrier integrity, with AECGM and PEx+ promoting a faster rate than DSI. After 30 hours in culture resistance in all media eventually stabilized indicating formation of a confluent monolayer with optimal barrier integrity. At this time the higher resistance observed in cells grown in PEx+ medium may also reflect their distinctive growth pattern. Unlike cells cultured in AECGM or DSI, which form typical “cobblestone” monolayers, cells in PEx+ do not exhibit contact inhibition and tend to grow in multiple layers. This results in a significantly higher cell density per unit area, contributing to increased resistance. Capacitance measurements revealed media-dependent differences., Capacitance measurements at high frequencies (≥32 kHz) decreased with time in all media but the rate of decline in PEx+ was markedly higher with baseline levels observed within 10 hours (**Fig. 5B, F-G**). In 2 donors AECGM also exhibited similar rapid decline in capacitance. These results suggest that AECGM and PEx+ promote more efficient cell adhesion and spreading than DSI. These media-dependent differences were further highlighted by plotting changes in resistance and capacitance at the optimal detection frequencies of 4 kHz and 64 kHz respectively (**Fig. 5C-H**). Specifically, in AECGM, resistance increased from 300 Ω to 1 kΩ within 5 hours, followed by a drop to ∼750 Ω at 12 hours. In DSI, resistance showed a steady increase from 300 Ω to ∼850 Ω over 12 hours, while in PEx+, there was a dramatic rise from 300 Ω to 4.3 kΩ within the first 12 hours (**Fig. 5C-D**). These increases in resistance were mirrored by similar patterns of capacitance decreases (**Fig. 5F-G**). The data shown in **Figure 5A**-**H** represent readings taken from one donor and is representative of the trends observed in two additional donors whose data is included in **Figure 5I** and available in the data supplement. Taken together, these ECIS data indicate that expansion in AECGM or PEx+ enables BCs to form tight junctions and reach confluency much faster than DSI. This strongly implicates pathways involved in TGF-β signaling which is a significant modulator of cell-matrix interactions.

**Figure 5:**
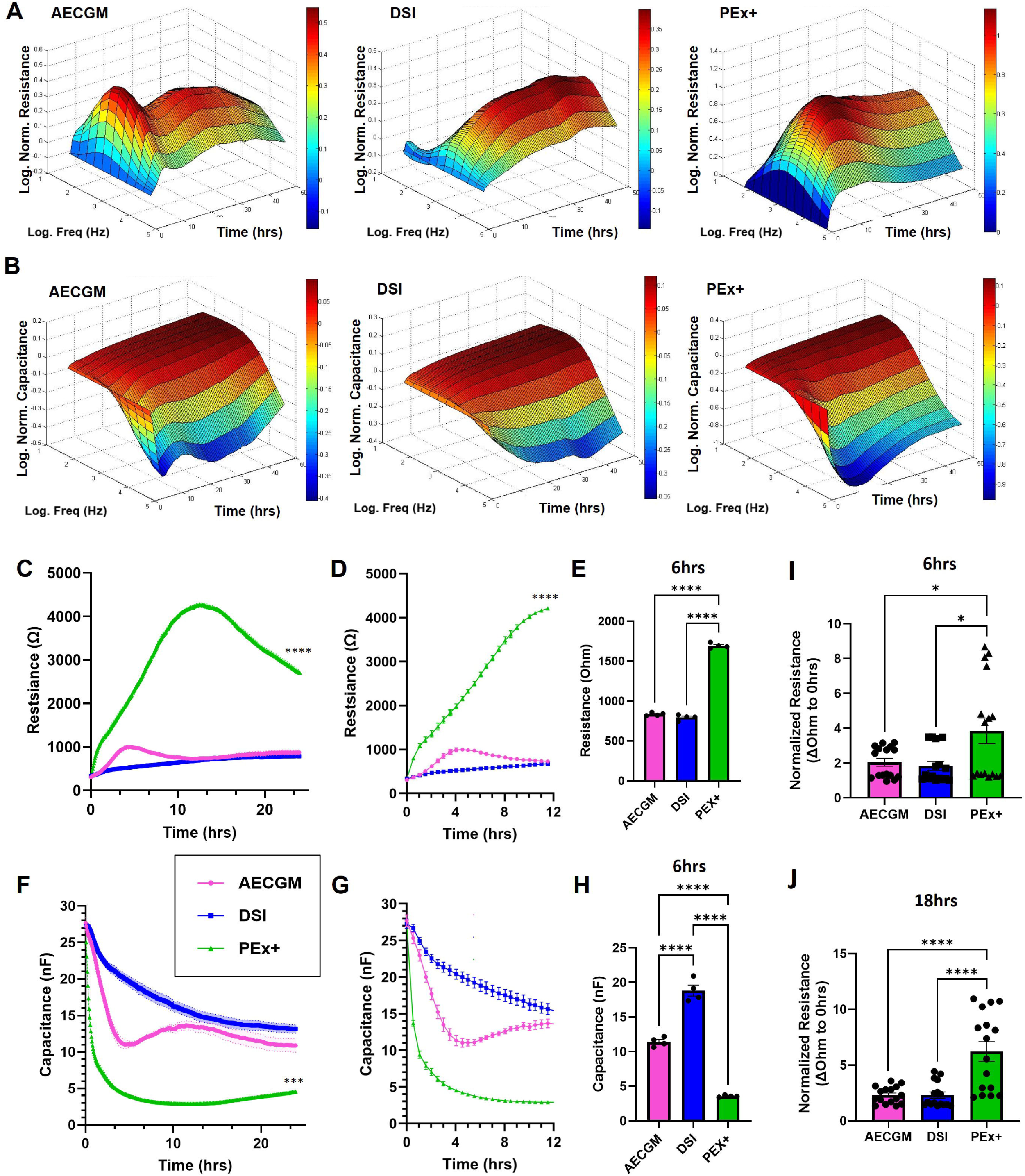
Culture Conditions Significantly Alter Biophysical Properties of BCs. ECIS measurements of BCs from a single donor cultured in the AECGM, DSI, or PEx+ media demonstrate marked differences in resistance (**A, C**) and capacitance **(B,F)** over 24 hours. Resistance at 4,000Hz shown over 24 hours **(C)** and 12 hours **(D)** for a single donor (n=4). (**E**) Measurement of resistance from one donor (n=4) after 6 hours. Capacitance at 64,000Hz shown over 24 (**F**) and 12 (**G**) hours. (**H**) Measurement of capacitance from one donor (n=4 technical replicates) after 6 hours. Resistance, normalized to 0-hour time point, at 4,000Hz from 3 independent donors after 6-hour (**I**) and 18-hour (**J**) incubation in the 3 media (N=3 donors, n=4 wells/donor). Mean±SEM. *p<0.05, ****p<0.0001.

### Culture-dependent alterations in BC niche interactions

To further understand the media-specific differences in gene regulation and biological processes we compared differential pathways enrichment, as calculated by IPA, that were observed between high (+) and low (-) passage gene expression changes within each indicated media type, and then compared those changes to the other media conditions. The comparison between time-influenced changes in AECGM and PEx+ (**Fig. 6A**), AECGM and DSI (**Fig. 6B**), and DSI and PEx+ (**Fig. 6C**) demonstrate that cell cycle and DNA replication related pathways were similarly changed in all medias (red and blue quadrants). Others including idiopathic pulmonary fibrosis (IPF), sirtuin signaling, ferroptosis, immune response and inflammation related pathways were differentially dependent on the media condition in which the cells were grown (**Fig. 6A-D**). The supporting data files are available in **Supplemental Tables S6-S8**. To highlight some notable changes in the RNAseq data we compared the normalized read counts of genes involved in cell-matrix interactions such as COL7A1, FN1, MMP-1, THBS1, and VIM and in cell surface adhesion and signaling such as FGFR3, ITGAV, ITGB6 and SERPINE1 in all 3 media across low and high passages (**Fig. 6E**). Many of these genes were higher in AECGM compared to DSI or PEx+ with levels of some such as COL7A1 and SERPINE1 demonstrating significant increase over time in culture (**Fig. 6E**). This was validated at the protein level. Fibronectin 1 and vimentin expression were higher in BCs expanded in AECGM compared to DSI or PEx+. Whereas fibronectin1 exhibited a significant increase in AECGM over time in culture, vimentin demonstrated a non-significant increase due largely to donor variability (**Fig. 6F**).

**Figure 6.**
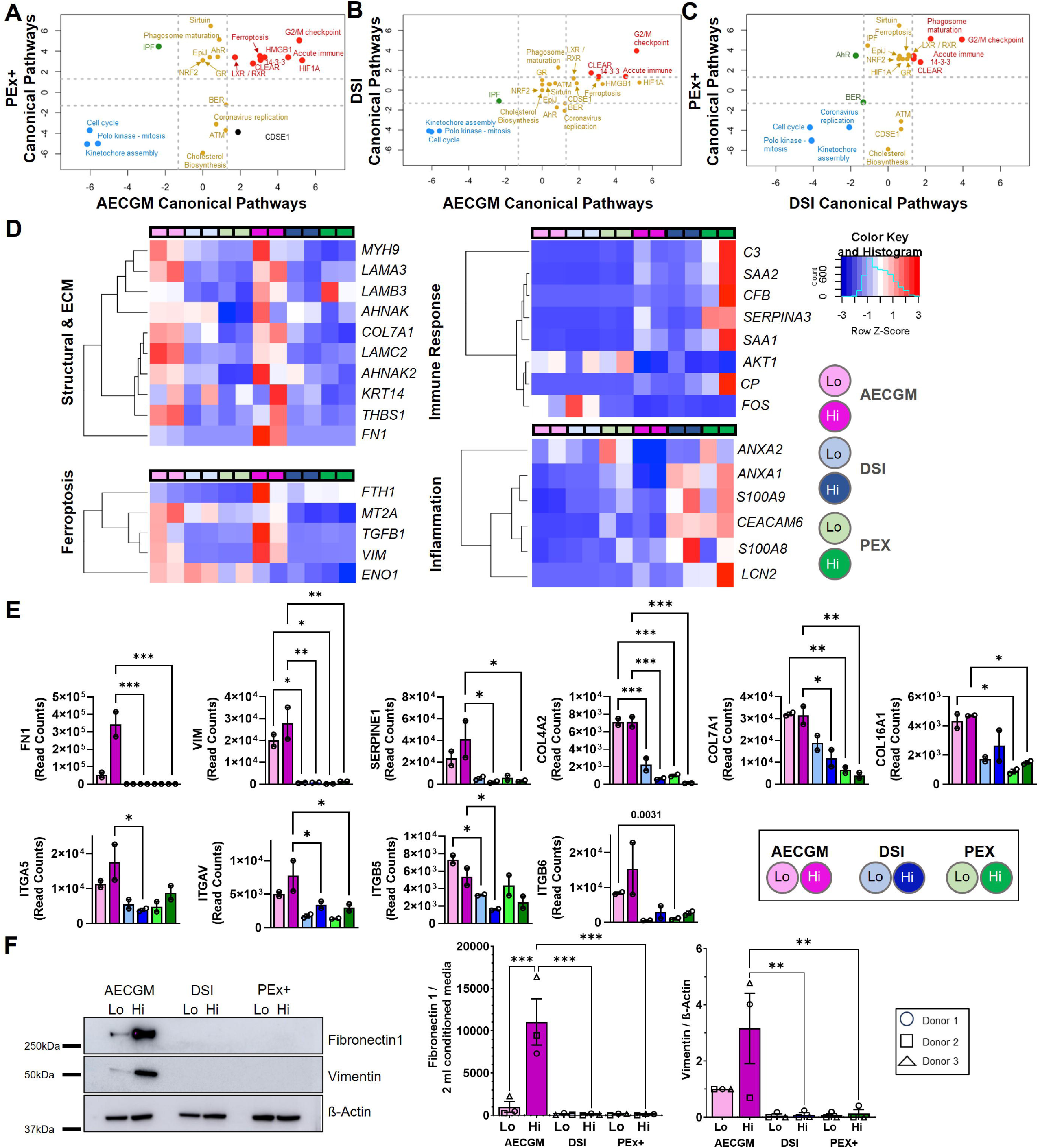
Altered Transcriptome Pathways by Long-Term Culture. Transcriptome analyses of most altered canonical pathway genes in cells cultured in AECGM and PEx+ (**A**), AECGM and DSI (**B**), and DSI and PEx+ (**C**). Blue=down regulated in both media, mustard=unchanged in the x axis media but significantly changed in the y axis media, green differentially regulated by media and red=upregulated in both media. (**D**) Heatmaps of genes reflecting the differential pathway regulation by the 3 media for specific genes selected to highlight differences in AECGM and PEx+ demonstrating changes in structural and ECM, immune response, and inflammation related genes. (**E**) RNAseq read counts of genes involved in cell-matrix interactions and cell surface adhesion and signaling for FN1, VIM, SEPRINE1, COL4A2, COL7A1, COL16A1, ITGA5, ITGAV, ITGB5 and ITGB6 in all 3 media across low and high passages. (**F**) Representative immunoblot of BCs from a single donor is shown. Densitometric quantitation of fibronectin 1 and vimentin from 3 donors normalized to conditioned media from 1 well and β-Actin, respectively. Data represents mean mean±SEM. *p<0.05, ** p<0.01, *** p<0.001, **** p<0.0001. Where significance is numerically stated this represents an unpaired student’s t-test.

Based on the observed increase in inflammation and immune response related gene expression in PEx+ and DSI cultures, we addressed the impact of AECGM, DSI and PEx+ media on BC inflammation. We collected conditioned medium from low or high passage BCs cultured in the three media and performed a multiplexed ELISA using Mesoscale Discovery (MSD) cytokine panels (**Fig. 7**). We found a significant increase in pro-inflammatory cytokine production from cells cultured in PEx+ at low passage, which persisted, albeit at reduced levels, at higher passages. The elevated pro-inflammatory cytokines included TNF-α, IL-1ß, IFN-g, and IL-6 (**Fig. 7A**). T cell regulatory cytokines such as IL-4, IL-13, IL-12p70, and IL-2 were also increased (**Fig. 7B**), along with the neutrophil chemoattractant IL-8 (**Fig. 7C**). No significant changes were observed for monocyte or dendritic cell chemokines, MCP-1, MCP-4, or MIP-1α. Additionally, PEx+ culture conditions were associated with increased levels of the anti-inflammatory cytokine IL-10 and IP-10, a CXCR3 ligand known to mediate proliferation and chemotaxis in bronchial epithelial cells (**Fig. 7D**). In contrast, cells cultured in DSI did not produce any of the pro-inflammatory cytokines or chemokines measured (**Fig. 6D** and **Fig. 7A-D**). However, extended culture in AECGM induced a specific pro-inflammatory signature, characterized by significant increases in the expression of pro-inflammatory genes such as IL-1β and IL-6, as well as the T cell modulator IL-2 (**Fig. 7 A-B**). These findings suggest that PEx+ culture promotes a broadly inflammatory microenvironment, which could have a substantial impact on both basal cell (BC) function and the BC niche.

**Figure 7.**
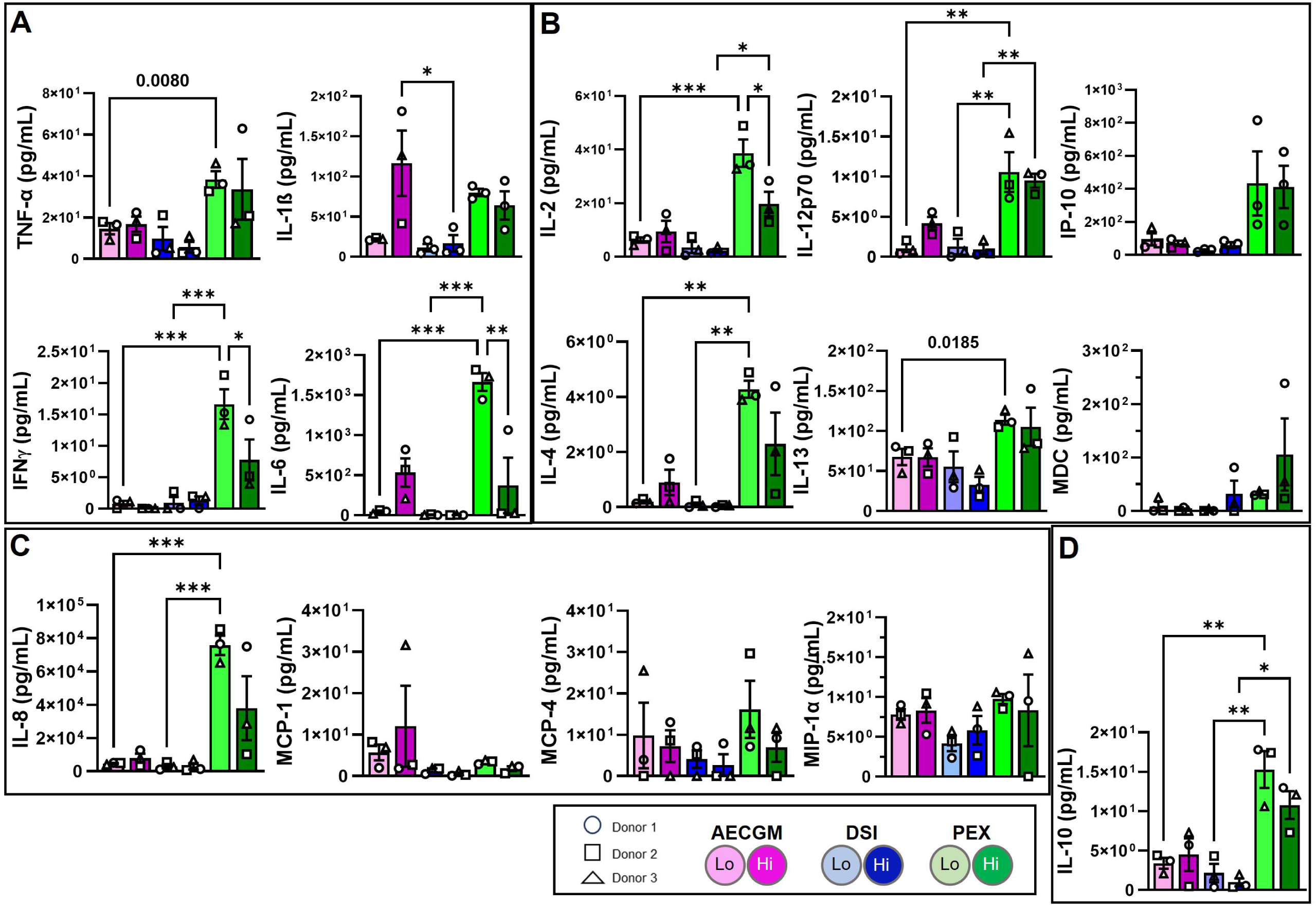
Media- and time-dependent alterations in inflammatory profile of BCs in culture. Secretion of cytokines and chemokines by BCs comparing low and high passage BCs expanded in AECGM, DSI or PEx+. **(A)** Pro-inflammatory cytokines: TNF-α, IL-1ß, IFN-γ, and IL-6. (**B**) T cell regulatory cytokines: IL-2, IL-12p70, IP-10, IL-4, IL-13, and MDC. **(C)** Neutrophil and monocyte chemoattractants: IL-8, MCP-1, MCP-4 and MIP-1α. (**D**) Anti-inflammatory cytokine IL-10. Data represents mean±SEM *p<0.05, ** p<0.01 *** p<0.001. Where significance is numerically stated this represents an unpaired student’s t-test.

## DISCUSSION

BCs experience significant transcriptomic, functional, and phenotypic changes when cultured for extended periods, limiting their usefulness in applications like autologous cell therapy. To address this, researchers have explored modifications to commercial media, particularly targeting the TGF-β superfamily with specific inhibitors. The BMP/TGF-β/SMAD signaling axis plays a key role in balancing BC quiescence and activation^36, 37^. Inhibiting dual SMAD activation, specifically in p63α+ BCs, has been shown to extend their proliferative capacity up to 80 population doublings^7^. Similar results have been achieved using irradiated 3T3 feeder layers, which degrade TGF-β, along with Rho kinase inhibition^17, 18^. While past studies mainly focused on BC growth and differentiation, their effects on other BC functions, such as matrix deposition and secretion of inflammatory mediators, remain underexplored, despite their importance in shaping the BC niche and regulating stemness.

Contrary to previous reports, we did not observe media-dependent changes in BC proliferation over time^7, 17, 18^. At low passage, BCs cultured in PEx+ exhibited a faster growth rate than those in AECGM or DSI, however, the higher resistance observed in ECIS experiments may also result from PEx+-cultured cells forming multiple layers, leading to higher cell density. Interestingly, cell cycle-promoting gene expression was similar across all media at low passage, suggesting equivalent proliferation rates. However, after 5-6 passages, cell cycle gene expression and p63α+ BC numbers declined significantly in all media, with a more pronounced reduction in PEx+, indicating that BCs in PEx+ exhaust faster than in AECGM or DSI. We did not co-stain with cell division markers, so we cannot rule out the presence of non-proliferative p63α+ BC subpopulations^14^. We limited culture expansion to about 16 population doublings, as BC growth rates declined by passage 5 in all media. This contrasts with Mou et al.’s findings of over 80 doublings with dual SMAD inhibition^7^, potentially due to differences in the matrix used for cell culture, collagen in our case versus laminin-rich 804G conditioned medium in their study^7, 38, 39^.

KRT5 is critical for maintaining the progenitor status of basal cells (BCs) and is increasingly linked to repair processes after injury^40, 41^. In idiopathic pulmonary fibrosis (IPF), KRT5+/p63+ BCs, usually found in the conducting airways, appear in distal airways within areas of bronchiolization, indicating that matrix remodeling or inflammation may drive KRT5+ cell migration and expression^42^. This is supported by evidence showing that TGF-β, AP-1 and NF-κB regulate KRT5 transcription^43, 44^. In our study, high-passage cells cultured in AECGM exhibited significantly higher TGF-β transcript levels compared to those in DSI or PEx+, potentially explaining the modestly higher KRT5 expression observed in these cells. However, post-transcriptional mechanisms might also play a role, as RNAseq data did not support transcriptional regulation of KRT5 in AECGM-cultured cells (**Fig. 3G**). Notably, robust KRT5 expression was also seen in low-passage cells cultured in DSI, which contains TGF-β inhibitors. Previous studies using various commercially available media to differentiate BCs at the ALI revealed Pneumacult ALI medium’s tendency to produce thicker epithelia with lower transepithelial resistance (reviewed in ^45^). Other factors, such as CBF, ciliated cell coverage, and MUC5AC and CFTR expression, varied across media. Our study differed by culturing BCs in different growth media (AECGM, DSI, or PEx+) before differentiation in Pneumacult ALI medium. We found that serial passaging, rather than the media itself, primarily affected differentiation into ciliated, MUC5AC+ secretory, and club cells, likely by reducing BC self-renewal.

Notably, we observed increased PLUNC expression in cells expanded in AECGM or PEx+. High-passage cells expressing PLUNC and CC10 exhibited diffuse intracellular staining and an enlarged phenotype, reminiscent of senescent cells. Given the reported effects of cytokines on goblet cell metaplasia^46^ and that senescent cells are associated with a secretory phenotype (SASP), including pro-inflammatory cytokines, growth factors and proteases^47, 48^, the elevated pro-inflammatory cytokines observed in p5 cells cultured in AECGM and PEx+ could contribute to cellular senescence^47, 49, 50^ and drive PLUNC expression. This is further supported by the fact that DSI, which suppressed inflammation, inhibited PLUNC expression in high-passage cells. However, the inflammatory profile in low passage PEx+ cultured cells suggests that inflammation alone may not fully explain the increased PLUNC expression Instead, PLUNC, itself may modulate epithelial cell inflammation. Recent observations by Ahmad et al demonstrate that intranasal administration of a PLUNC peptidomimetic inhibited bacterial burden and mortality in a mouse model of acute lung injury^51^. By targeting the calcium-release activated calcium channel protein 1 (Orai-1), this mimetic inhibited NFAT-mediated inflammatory gene expression. Although the anti-inflammatory effects of this mimetic were largely observed in alveolar macrophages, its effects on airway epithelial cells cannot be excluded as these cells express the *ORAI1* gene^52^. If true and given its known antimicrobial properties^53^, increased PLUNC expression in serially passaged BCs in AECGM and PEx+ may indeed be a protective mechanism to guard against excessive inflammation-related damage and BC exhaustion.

The ECM is crucial for BC function, providing structural support and regulating growth factor signaling^1^. Multiple ECM proteins are known to bind growth factors modulating their stability, availability, and duration of signaling^54–56^. For instance, by sequestering IGFBP3 which binds IGF, collagen regulates IGF-dependent cellular processes such as proliferation^56^. Our RNAseq data showed that AECGM cultured cells exhibited higher expression of ECM components such as collagen7A1, fibronectin1, and TGF-β1, indicating enhanced matrix-mediated regulation of BC functions in this medium. Moreover, these cells showed increased expression of integrin genes (ITGβ6 and ITGαV), suggesting improved migratory and adhesive properties^57, 58^. SERPINE1, which modulates cell-matrix interactions, was also enriched in AECGM cultured cells, potentially affecting BC adhesion and migration through its interaction with other matrix components^59, 60^. In addition, Serpine1 mRNA, independent of the encoded SERPINE1 protein, promotes cell migration by sequestering miRNA^61^. Given that SERPINE1 is activated by TGF-ß1^62^ and αV integrins activate TGF-ß1^63, 64^, the effects of TGF-ß1 signaling on BC-matrix interactions cannot be ruled out. These effects may be especially relevant in cell-based therapeutic approaches to treat chronic lung diseases characterized by dysregulated matrix remodeling and lung repair. Our data would predict that cells expanded in AECGM, when transplanted in patients with chronic lung disease, would exhibit greater ability to regulate fundamental processes in lung repair such as migration to sites of injury, adhesion, and self-renewal.

Inflammatory mediators play a critical role in BC function and airway regeneration (reviewed in ^65, 66^). Our data revealed that media conditions significantly influence the inflammatory niche of BCs, with PEx+ cultured cells exhibiting a broad inflammatory profile, regardless of culture duration, indicating activation of multiple stress-regulated pathways^67–71^. This notion is supported by our observation of enrichment of NRF2 oxidative stress pathway in BCs expanded in PEx+. The broad inflammatory response to PEx+ medium occurred despite increased secretion of the anti-inflammatory cytokine IL-10, suggesting a compensatory mechanism against inflammation. Importantly, these cytokine differences were independent of hydrocortisone levels, which were consistent across all media. The proprietary composition of PEx+, unlike AECGM and DSI, complicates definitive conclusions about its effects. In contrast to PEx+, AECGM promoted a more targeted pro-inflammatory response whereas DSI suppressed inflammation altogether, thus implicating the TGF-β1/BMP pathway in BC mediated inflammatory processes. Airway epithelial cells are inherently programmed to initiate inflammatory gene expression in response to injury or infection, facilitating airway regeneration^72–75^. The IL-1β-TNF-α-NF-κB signaling axis, for example, supports alveolar repair following influenza infection^76^. Similarly, injury or protease activity can stimulate IL-6 release by HBECs, enhancing multiciliogenesis at the expense of secretory cells^77, 78^. IL-13 plays a contrasting role by increasing mucin-secreting goblet cells while reducing ciliated cells in asthma, where TH2-driven goblet cell hyperplasia occurs^79^. TH_2_ cytokines, including IL-4 and IL-13, also enhance tracheosphere size^46^ and promote epithelial repair through mechanisms involving M2 macrophage polarization and CCR2+ monocyte recruitment^80, 81^. Our data reveal significant effects of media conditions on the constitutive expression of these cytokines, with levels often matching or exceeding those found in BALF from lung disease patients^82–84^. Based on these previous reports, it is expected that the distinct inflammatory profile promoted by PEx+ is likely to impact BC function in a dramatically different manner to AECGM or DSI. These results highlight the importance of culture conditions on BC-inflammatory niche interactions which are critical in shaping BC behavior.

In conclusion, despite the many *in vitro* models of airway disease, the impact of culture conditions on inflammatory responses of airway epithelial cells has not been systematically examined. The media-dependent effects on inflammation discovered in our study underscore the importance of considering culture conditions when interpreting *in vitro* BC studies or predicting BC behavior following transplantation.

## DATA AVAILABILITY

All next-generation sequencing data utilized in this study is publicly available through the National Center for Biotechnology Innovation Gene Expression Omnibus (NCBI-GEO) as record GSE224243.

## SUPPLEMENTAL MATERIAL

An Online Data Supplement is available at doi: 10.6084/m9.figshare.26813173. This includes:

- Supplemental Tables S1-S7 doi: 10.6084/m9.figshare.26793058
- Supplemental Figures S1 doi: 10.6084/m9.figshare.26793052
- Supporting Raw data files doi: 10.6084/m9.figshare.26792782

## AUTHOR CONTRIBUTIONS

SM and CM: Conceived and designed research, performed experiments, analyzed data, interpreted results of experiments, prepared figures, drafted manuscript, edited and revised manuscript, approved final version of manuscript. DS, AC and SB: performed experiments, analyzed data, interpreted results of experiments, and approved final version of manuscript. BAC and LKG: performed experiments, analyzed data, interpreted results of experiments, edited, and revised manuscript, and approved final version of manuscript. AN: analyzed data. ALR: conceived and designed research, performed experiments, analyzed data, interpreted results of experiments, prepared figures, drafted manuscript, edited and revised manuscript, approved final version of manuscript and was responsible for funding acquisition.

