## Supplementary material for "Culture Conditions Differentially Regulate the Inflammatory Niche and Cellular Phenotype of Tracheo-Bronchial Basal Stem Cells": Online Data Supplement

RUNNING HEAD: Regulation of Airway Basal Cell Phenotype

### Differential Regulation of Tracheo-Bronchial Basal Cell Phenotype By Culture Conditions

**Supplemental Tables**

All supplemental Tables can be found on Fig Share at doi:10.6084/m9.figshare.26793058.

**Supplemental Table S1:** Airway basal cell donor information for cells used in this study.
**Supplemental Table S2.** Detailed description of medias used in this study.
**Supplemental Table S3:** Antibodies used in this study.
**Supplemental Table S4.** Read count per transcript, adjusted raw sequencing reads.
**Supplemental Table S5:** IPA pathways associated with PEx+ and AECGM media expansion.
**Supplemental Table S6:** IPA pathways associated with DSI and AECGM media expansion.
**Supplemental Table S7:** IPA pathways associated with PEx+ and DSI media expansion.

#### Supplemental Figures

All Supplemental Figures can be found on Fig Share at doi: 10.6084/m9.figshare.26793052

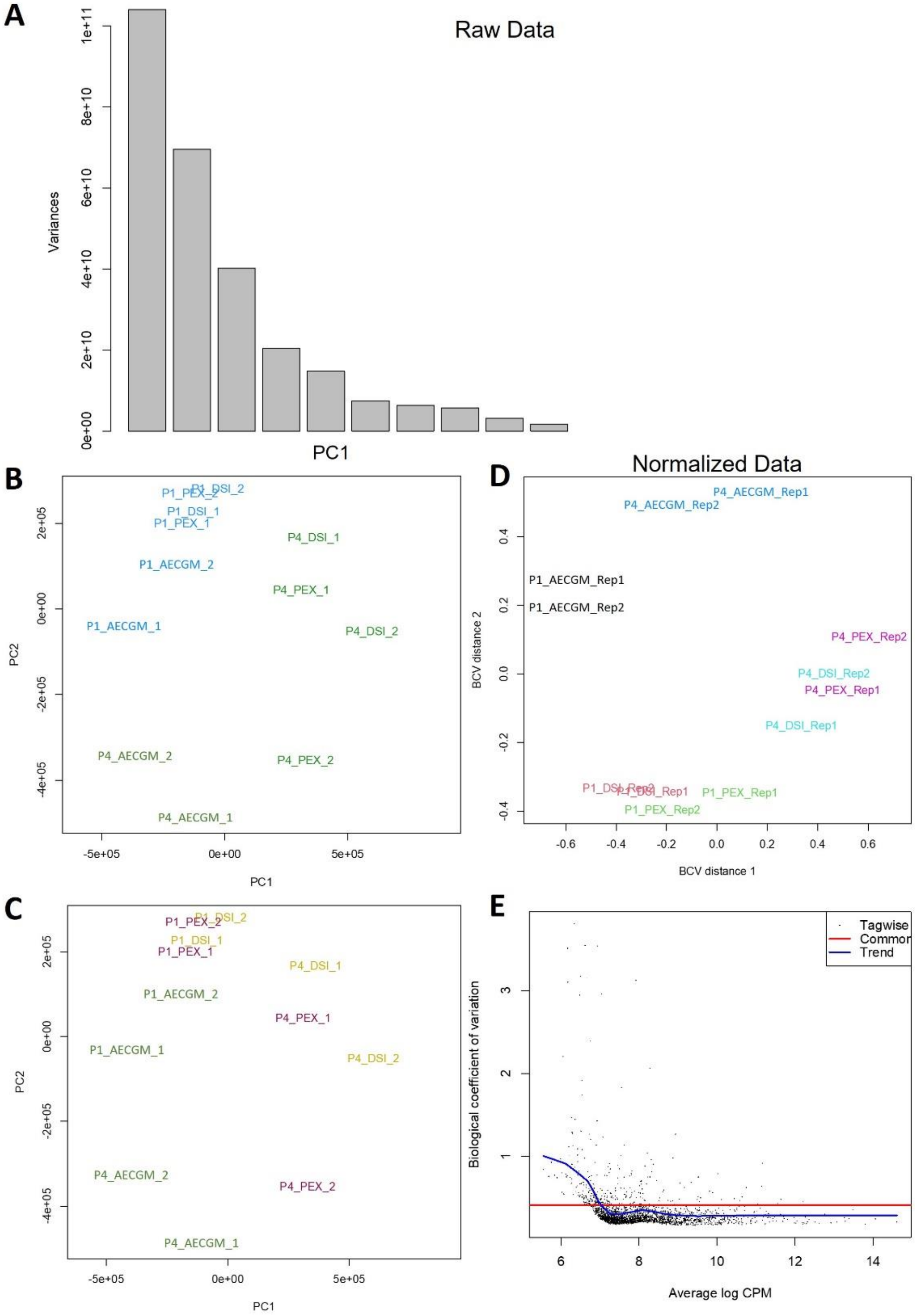


**Supplemental Figure S1: Quality Control metrics on Bulk RNAseq BC media treatment and time in culture dataset.** (A) Bar plot of principal components contributing to variation within the data and their relative contribution. Y-axis = variance contribution to the overall dataset. (B) Principal component analysis of Raw data (pre-normalization) colored by time in culture. X-axis = Principal Component 1, Y-axis = Principal Component 2. Blue = low passage cells, green = high passage cells.(C) Principal Component analysis of raw data (pre-normalization) colored by treatment. Blue = AECGM/BEGM media, mustard = DSI media, red = PEx+ media. (D) Principal Component analysis of normalized bulk RNAseq data. Black = AECGM/BEGM at low passage, blue = AECGM/BEGM at high passage, red = DSI media at low passage, teal = DSI media at high passage, green = PEx+ media at low passage, purple = PEx+ media at high passage. (E) Biologic coefficient of variance across the entire bulk RNAseq dataset. Red line = common dispersion, blue line = common dispersion trend, black points = gene dispersions across the dataset. X-axis = average expression in log counts per million (log CPM) across the normalized dataset. Y-axis = biological coefficient of variation for each gene across the dataset.
